# Unraveling rice tolerance mechanisms against *Schizotetranychus oryzae* mite infestation

**DOI:** 10.1101/281733

**Authors:** Giseli Bufon, Édina Aparecida dos Reis Blasi, Angie Geraldine Sierra Rativa, Thainá Inês Lamb, Rodrigo Gastmann, Janete Mariza Adamski, Joséli Schwambach, Felipe Klein Ricachenevsky, Angelo Schuabb Heringer, Vanildo Silveira, Mara Cristina Barbosa Lopes, Raul Antonio Sperotto

## Abstract

Infestation of *Schizotetranychus oryzae* (Acari: Tetranychidae) causes great losses in rice productivity. Infestation in Puitá INTA-CL cultivar reduced the number of seeds/plant, percentage of full seeds, 1,000 seeds weight, and seed length, whereas infestation in IRGA423 increased 1,000 seeds weight and seed length. Reduction in seed weight/plant caused by infestation was higher in Puitá INTA-CL than IRGA423. Thus, Puitá INTA-CL was established as susceptible, and IRGA423 as tolerant to *S. oryzae* infestation. Photosynthetic parameters were less affected by infestation in IRGA423 than in Puitá INTA-CL. Infestation also caused accumulation of H_2_O_2_, decreased cell membrane integrity and accelerated senescence in leaves of Puitá INTA-CL, while leaves of IRGA423 presented higher levels of phenolics compounds. Using proteomic analysis, we identified proteins related to plant defense, such as jasmonate synthesis, and related to other mechanisms of tolerance such as oxidative stress, photosynthesis, and DNA structure maintenance, more abundant in IRGA423 after seven days of infestation. We detected higher levels of silicon (as amorphous silica cells) in leaves of infested IRGA423 plants compared to Puitá INTA-CL, an element previously linked to plant defense. Our data shows that IRGA423 presents tolerance to *S. oryzae* infestation, and that multiple mechanisms might be employed by this cultivar.

**Highlight:** This is the first report evaluating the defense responses (tolerance and susceptiblity) of two contrasting rice cultivars to *Schizotetranychus oryzae* mite infestation.

## Introduction

Rice is one of the most important sources for global food security and socioeconomic stability (FAO, 2017). Research directed to this crop are important for the development of technologies that increase productivity and assist farmers who depend on it for subsistence, as is the case in several developing countries such as Brazil (Zeigler and Barclay, 2008), which is the ninth largest rice producer and the main producer outside Asia (FAO, 2017). In the last years Brazil produced around 10 million tons of rice, with Rio Grande do Sul (RS) state accounting for approximately 70% of this amount. However, monoculture and intensive use of fertilizers benefit the appearance of pest arthropods, which are the main competitors of humans for the resources generated by agriculture (Oerke and Dehne, 2004).

Interactions between plants and insect herbivores are important determinants of plant productivity in managed and natural vegetation. In response to attack, plants have evolved a range of defenses to reduce the threat of injury and seed set. Crop losses from damage caused by arthropod pests can exceed 15% annually (Mitchell *et al.,* 2016). In order to quantify the pest resistance of the cultivars, the best tool does not seem to be the increase of the arthropod population, but the measurement of the damages caused to the plants, since the reduction of the leaf damage is followed by an increase in yield and quality of the grain, and these are the ultimate objectives of most crop breeding programs (Smith, 2005). Thus, the plant resistance/tolerance to arthropods is the sum of genetically inherited traits that result in an adapted species that suffers less damage compared to susceptible ones. These resistance/tolerance qualities should be measured on a relative scale by comparing levels of damage and productivity with susceptible plants that are severely damaged when exposed to similar experimental conditions (Smith, 2005). Plant tolerance to arthropods has been indicated as a category of resistance. However, very little is known about the mechanism of tolerance to arthropods.

Tolerance is distinctive in terms of the plant’s ability to withstand or recover from herbivore injury through growth and compensatory physiological processes (Koch *et al.,* 2016). Since plant tolerance involves compensatory behavior, the plant is able to bear a large number of herbivores without interfering with the insect pest’s physiology or behavior. Some studies observed that tolerant plants can compensate photosynthetically by avoiding feedback inhibition and impaired electron flow through photosystem II that occurs as a result of insect feeding. Similarly, the up-regulation of peroxidases and other oxidative enzymes during insect feeding, together with elevated levels of phytohormones, can play an important role in plant tolerance to insect pests (Koch *et al.,* 2016).

Phytophagous mites (Acari) comprise a diverse group of arthropods with several species that are pests in crop plants. Within this group, the spider mites family Tetranychidae are of special interest since they cover a broad host-plant range and can develop into devastating outbreaks (Van Leeuwen *et al.,* 2015). Adult spider mites feed from leaves by piercing mesophyll cells with their stylets, and sucking the cell content (Villarroel *et al.,* 2016). Such feeding behavior can severely damage leaf tissues (Bensoussan *et al.,* 2016). To control such damage there is an indiscriminate use of acaricides. However, mites of the Tetranychidade family have been reported for developing resistance to various acaricides (Osakabe *et al.,* 2016). Among the mites from the Tetranychidae family found in rice crops that cause economic damage is *Schizotetranychus oryzae* Rossi de Simons, which has been reported in several South American countries, and generates damages in irrigated rice fields (Ferla *et al.,* 2013), which may become increasingly frequent due to global warming (Reddy, 2015). To date, there is little information about the damage and economic lost caused by *S. oryzae* infestation in rice crops, and the available information is usually related to visual effects of the plant. Recently, our group described differentially abundant proteins in rice leaves early infested (Buffon *et al.,* 2016) and late-infested (Blasi *et al.,* 2017) by *S. oryzae*, along with the physiological changes induced by such different mite populations. However, the molecular and physiological changes caused by *S. oryzae* infestation in contrastant resistant/tolerant and susceptible rice cultivars have not yet been elucidated.

Even though *S. oryzae* being the phytophagous mite most commonly found in rice cultivation in the RS state (Ferla *et al.,* 2013), field observations show that some rice cultivars present different levels of infestation, suggesting a possible resistance mechanism. Therefore, we evaluated different rice cultivars commonly cultivated in different regions of RS state aiming to identify different rice responses to *S. oryzae* infestation, in order to understand the molecular and physiological mechanisms behind resistance/tolerance and susceptibility to this mite. Our results may be useful for future breeding programs aiming at resistance/tolerance to phytophagous mite *S. oryzae* infestation.

## Material and methods

### Plant growth conditions and mite infestation

Seeds of rice (*Oryza sativa* L. ssp. *indica*) from IRGA 426, BRS Atalanta, Puitá INTA-CL, IRGA 424, BRS 7 Taim, IRGA 410, and IRGA 423 cultivars were surface sterilized and germinated for four days in an incubator (28°C) on paper soaked with distilled water. After germination, plantlets were transferred to vermiculite/soil mixture (1:3) for additional 14 days in greenhouse conditions, and then transferred to plastic buckets containing soil and water. Plastic buckets containing rice plants highly infested by *S. oryzae* were kindly provided by Instituto Rio-Grandense do Arroz (IRGA, Cachoeirinha, RS), and were used to infest rice plants in our experiment. Fifty plants (V7-9 stage, according to Counce *et al.,* 2000) of each cultivar (five plants per bucket) were infested by proximity with the bucket containing the highly infested plants placed in the center of the other buckets. For greater homogeneity of infestation and contact, buckets of each cultivar were rotated at a 90° angle counterclockwise every two days. Fifty plants of each cultivar were cultivated without infestation (control condition).

The level of damage caused by *S. oryzae* in the tested rice plants was analyzed during the average period of three months, until the plants reach its final stage of reproductive development (panicle maturity, R9 stage; Counce *et al.,* 2000). Cultivars that maintained the highest integrity of the leaves during infestation were characterized as possible resistant/tolerant to *S. oryzae*. Evaluation of damage in the abaxial and adaxial faces of leaves was based on a classification of four levels of infestation: control condition, without any sign of infestation; early infested (EI) leaves, 10 to 20% of damaged leaf area, average of 168 hs of exposure to the mite; intermediate infested (II) leaves, 40 to 50% of damaged leaf area, average of 360 hs; and late infested (LI) leaves, more than 80% of damaged leaf area, average of 720 hs, according to Fig. S1.

### Plant height and tiller number

Plant height and tiller number were evaluated during the vegetative stage (V7-9, before being infested) and during the last reproductive stage (R9, control and infested plants).

### Chlorophyll a fluorescence transients

The chlorophyll a fluorescence transient was measured on the third upper leaves of control and infested plants of the cultivars Puitá INTA-CL, BRS 7 Taim and IRGA 423 in three different exposure times: early infestation (EI), intermediate infestation (II), and late infestation (LI), using a portable fluorometer (OS30p, Optisciences, UK). Before the measurements, plants were dark adapted for 20 minutes and the fluorescence intensity was measured by applying a saturating pulse of 3,000 μmol photons m^−2^ s^−1^ and the resulting fluorescence of the chlorophyll a measured from 0 to 1 s, OJIP curve (Strasser *et al.,* 2000). This data was used to calculate parameters of the JIP Test (Strasser *et al.,* 2000; Tsimilli-Michael and Strasser, 2008).

### Seed analysis

Seeds from Puitá INTA-CL and IRGA 423 cultivars were collected in R9 stage and the following agronomical parameters were evaluated: number of seeds (empty + full) per plant, percentage of full seeds, weight of 1,000 full seeds, and seed length. Yield reduction caused by *S. oryzae* infestation was calculated using the following equation for each cultivar and each condition (control and infested): number of seeds (empty + full) per plant X percentage of full seeds X weight of one seed = seed weight per plant. The seed weight per plant of the infested condition was divided by the seed weight per plant of the control condition, showing an estimative of yield loss percentage in each cultivar caused by *S. oryzae* infestation.

### In situ histochemical localization of *H*_*2*_*O*_*2*_ and loss of plasma membrane integrity

*In situ* accumulation of H_2_O_2_ in control and late infested leaves of Puitá INTA-CL and IRGA 423 cultivars was detected by histochemical staining with diaminobenzidine (DAB), according to Shi *et al.* (2010), with minor modifications. For H_2_O_2_ localization, leaves were immersed in DAB solution (1 mg ml^−1^, pH 3.8) in 10 mM phosphate buffer (pH 7.8), and incubated at room temperature for 8 h in the light until brown spots were visible, which are derived from the reaction of DAB with H_2_O_2_. Leaves were bleached in boiling concentrated ethanol to visualize the blue and brown spots, and kept in 70% ethanol for photo documentation with a digital camera coupled to a stereomicroscope. To determine changes in cell viability (indicative of cell death), another set of control and late infested leaves were immersed for 5 h in a 0.25% (w/v) aqueous solution of Evans Blue (Romero-Puertas *et al.,* 2004). Leaves were discolored in boiling concentrated ethanol to develop the blue precipitates, which were photo documented with a digital camera coupled to a stereomicroscope.

### Phenolic compounds

Phenolic compounds were quantified according to Fett-Neto *et al.* (1992), with minor modifications. Approximately 50 mg of control and early infested leaves from Puitá INTA-CL and IRGA 423 cultivars were pulverized in liquid nitrogen, extracted in 2 ml 0.1 N HCl and submitted to sonication in a water bath for 30 min. The extracts were centrifuged at 9,000 rpm for 10 min at 4°C. The supernatant was collected and the pellet was re-extracted. The supernatants were pooled and the final volume was completed to 1,5 ml with 0.1 N HCl. For quantification, 300 μl of 20% (w/v) Na_2_CO_3_ and 150 μl of Folin-Ciocalteu reagent were added, mixed and then incubated at 100°C for 1 min. Absorbance was read at 750 nm. The standard curve was established with Gallic Acid in 0.1 N HCl.

### Microscope observation of amorphous silica cells and Silicon quantification

Control and late infested leaves from Puitá INTA-CL and IRGA423 cultivars were used in observation of amorphous silica cells and SiO_2_ quantification. Morphology of silica cells on the leaf surfaces (abaxial and adaxial faces) was observed using scanning electron microscopy (SEM). A fresh specimen (0.3-0.5 cm in length) part of the reciprocal fourth leaf was sampled and wiped with tissue paper to remove moisture. The leaf segment was fixed and coated with metal and then loaded onto the SEM. Pictures (at 700 × magnification) were obtained to illustrate the differences in amorphous silica cells of rice leaves.

### Protein extraction and quantification

Three biological samples (250 mg of fresh matter) of control and early infested leaves from Puitá INTA-CL and IRGA 423 cultivars, each containing three leaves from three different plants, were subjected to protein extraction using Plant Total Protein Extraction Kit (Sigma-Aldrich). The protein concentration was measured using 2-D Quant Kit (GE Healthcare, Piscataway, NJ, USA).

### Protein digestion

For protein digestion, three biological replicates of 100 μg of proteins were used. Before the trypsin digestion step, protein samples were precipitated using the methanol/chloroform methodology to remove any detergent from samples (Nanjo *et al.,* 2012). Then, samples were resuspended in Urea 7 M and Thiourea 2 M buffer, and desalted on Amicon Ultra-0.5 3 kDa centrifugal filters (Merck Millipore, Germany). Filters were filled to maximum capacity with buffers and centrifuged at 15,000 g for 10 min at 20°C. The washes were performed twice with Urea 8 M and then twice with 50 mM ammonium bicarbonate (Sigma-Aldrich) pH 8.5, remaining approximately 50 μL per sample after the last wash. The methodology used for protein digestion was as previously described (Calderan-Rodrigues *et al.,* 2014). For each sample, 25 μL of 0.2% (v/v) RapiGest® (Waters, Milford, CT, USA) was added, and samples were briefly vortexed and incubated in an Eppendorf Thermomixer® at 80°C for 15 min. Then, 2.5 μL of 100 mM DTT (GE Healthcare) was added, and the tubes were vortexed and incubated at 60°C for 30 min under agitation. Next, 2.5 μL of 300 mM iodoacetamide (GE Healthcare) was added, and the samples were vortexed and then incubated in the dark for 30 min at room temperature. The digestion was performed by adding 20 μL of trypsin solution (50 ng/μL; V5111, Promega, Madison, WI, USA) prepared in 50 mM ammonium bicarbonate, and samples were incubated at 37°C during 15 hs. For RapiGest® precipitation and trypsin activity inhibition, 10 μL of 5% (v/v) trifluoroacetic acid (TFA, Sigma-Aldrich) was added and incubated at 37°C for 30 min, followed by a centrifugation step of 20 min at 16,000 g. Samples were transferred to Total Recovery Vials (Waters).

### Mass spectrometry analysis

A nanoAcquity UPLC connected to a Synapt G2-Si HDMS mass spectrometer (Waters, Manchester, UK) was used for ESI-LC-MS/MS analysis. Fist was performed a chromatography step by injecting 1 μL of digested samples (500 ng/μL) for normalization to relative quantification of proteins. To ensure standardized molar values for all conditions, normalization among samples was based on stoichiometric measurements of total ion counts of MS^E^ scouting runs prior to analyses using the ProteinLynx Global SERVER v. 3.0 program (PLGS; Waters). Runs consisted of three biological replicates. During separation, samples were loaded onto the nanoAcquity UPLC 5 μm C18 trap column (180 μm × 20 mm) at 5 μL/min during 3 min and then onto the nanoAcquity HSS T3 1.8 μm analytical reversed phase column (75 μm × 150 mm) at 400 nL/min, with a column temperature of 45°C. For peptide elution, a binary gradient was used, with mobile phase A consisting of water (Tedia, Fairfield, Ohio, USA) and 0.1% formic acid (Sigma-Aldrich), and mobile phase B consisting of acetonitrile (Sigma-Aldrich) and 0.1% formic acid. Gradient elution started at 7% B, then ramped from 7% B to 40% B up to 91.12 min, and from 40% B to 99.9% B until 92.72 min, being maintained at 99.9% until 106.00 min, then decreasing to 7% B until 106.1 min and kept 7% B until the end of experiment at 120.00 min. Mass spectrometry was performed in positive and resolution mode (V mode), 35,000 FWHM, with ion mobility, and in data-independent acquisition (DIA) mode; ion mobility separation (HDMS^E^) using IMS wave velocity of 600 m/s, and helium and IMS gas flow of 180 and 90 mL/min respectively; the transfer collision energy ramped from 19 V to 55 V in high-energy mode; cone and capillary voltages of 30 V and 2750 V, respectively; and a source temperature of 70°C. In TOF parameters, the scan time was set to 0.5 s in continuum mode with a mass range of 50 to 2,000 Da. The human [Glu1]-fibrinopeptide B (Sigma-Aldrich) at 100 fmol/μL was used as an external calibrant and lock mass acquisition was performed every 30 s. Mass spectra acquisition was performed by MassLynx v4.0 software.

### Bioinformatics

Spectra processing and database searching conditions were performed by Progenesis QI for Proteomics Software V.2.0 (Nonlinear Dynamics, Newcastle, UK). The analysis used the following parameters: Apex3D of 150 counts for low energy threshold, 50 counts for elevated energy threshold, and 750 counts for intensity threshold; one missed cleavage, minimum fragment ion per peptide equal to two, minimum fragment ion per protein equal to five, minimum peptide per protein equal to two, fixed modifications of carbamidomethyl (C) and variable modifications of oxidation (M) and phosphoryl (STY), and a default false discovery rate (FDR) value at a 1% maximum, peptide score greater than four, and maximum mass errors of 10 ppm. The analysis used the *Oryza sativa* protein databank from Phytozome (https://phytozome.jgi.doe.gov/). Label-free relative quantitative analyses were performed based on the ratio of protein ion counts among contrasting samples. After data processing and to ensure the quality of results, only proteins present or absent (for unique proteins) in three out of three runs were accepted and submitted to differentially abundance analysis. Proteins were considered to be up-regulated if the fold change (FC) was greater than 1.5 and down-regulated if the FC was less than 0.6667, and both with significantly P-value ANOVA (P < 0.05). The Blast2GO tool (http://www.blast2go.org) was used to identify proteins with known Gene Ontology annotations (Conesa *et al.,* 2005). Detected proteins were also analyzed using the B2G Kegg maps (Ashburner *et al.,* 2000).

### RNA extraction and cDNA synthesis

Total RNA was extracted from control and early infested rice leaves of Puitá INTA-CL and IRGA 423 cultivars using NucleoSpin RNA Plant (Macherey-Nagel). First-strand cDNA synthesis was performed using the SMART PCR cDNA Synthesis Kit (Clontech Laboratories, Mountain View, CA, USA) with reverse transcriptase (M-MLV, Invitrogen, Carlsbad, CA, USA) and 2 μg of RNA quantified using Qubit RNA Assay Kit (Invitrogen) and Qubit 2.0 Fluorometer.

### Quantitative RT-PCR and data analysis

RT-qPCRs were carried out in a StepOne Real-Time Cycler (Applied Biosystems). All primers (listed in Table S1) were designed to amplify 100-150 bp of the 3’-UTR of the genes and to have similar Tm values (60 ± 2°C). Reaction settings were composed of an initial denaturation step of 5 min at 94°C, followed by 40 cycles of 10 s at 94°C, 15 s at 60°C, 15 s at 72°C and 35 s at 60°C (fluorescence data collection); samples were held for 2 min at 40°C for annealing of the amplified products and then heated from 55 to 99°C with a ramp of 0.1°C/s to produce the denaturing curve of the amplified products. RT-qPCRs were carried out in 20 μl final volume composed of 10 μl of each reverse transcription sample diluted 100 times, 2 μl of 10X PCR buffer, 1.2 μl of 50 mM MgCl_2_, 0.1 μl of 5 mM dNTPs, 0.4 μl of 10 μM primer pairs, 4.25 μl of water, 2.0 μl of SYBR green (1:10,000, Molecular Probe), and 0.05 μl of Platinum Taq DNA polymerase (5 U/μl, Invitrogen, Carlsbad, CA, USA). Gene expression was evaluated using a modified 2^−ΔCT^ method (Schmittgen and Livak, 2008), which takes into account the PCR efficiencies of each primer pair (Relative expression TESTED GENE/CONTROL GENE = (PCR_eff_ CG)^Ct CG^ / (PCR_eff_ TG)^Ct TG^). *OsUBQ5* gene expression was used as internal control to normalize the relative expression of tested genes (Jain *et al.,* 2006). Each data point corresponds to three biological and four technical replicate samples. The expression of a senescence marker gene (*Staygreen* gene, *OsSGR*, a chloroplast protein which regulates chlorophyll degradation by inducing LHCII disassembly through direct interaction; Park *et al.,* 2007) was also analyzed in control and early/intermediate infested leaves.

### Statistical analysis (except for proteomic/bioinformatic analyzes)

Data were analyzed using the Student’s *t* test (P ≤ 0.05, 0.01, and 0.001) or One-Way ANOVA followed by Tukey test, using SPSS Base 21.0 for Windows (SPSS Inc., USA).

## Results

### Different physiological responses of rice cultivars to S. oryzae infestation

The first screening of rice responses to *S. oryzae* infestation showed that all tested cultivars present similar pattern of infestation kinetics (Fig. S2). After five weeks, high infestation levels were detected in all cultivars. Therefore, none of the tested cultivars seems to be resistant to *S. oryzae* infestation. On the other hand, plant height and tiller number were differentially affected by mite infestation (Fig. 1). *S. oryzae* affected the plant growth of IRGA 426, Puitá INTA-CL and IRGA 410 cultivars, and also the tillering in BRS Atalanta, Puitá INTA-CL and IRGA 423 cultivars. Even though we were not able to find any sign of resistance in these cultivars, we decided to further characterize the response to *S. oryzae* of three cultivars that showed different responses to mite infestation. Therefore, we selected Puitá INTA-CL, BRS7 Taim and IRGA 423 cultivars to further analysis.

**Fig. 1.**
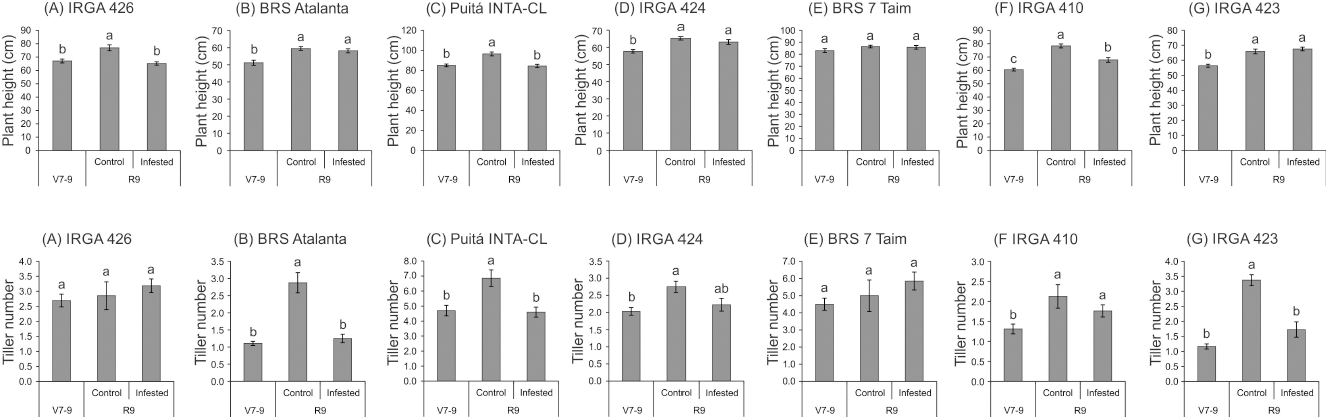
Plant height (cm) and tiller number of the seven tested cultivars at the vegetative stage (no infestation, V7-9) and full maturity stage (control or infested conditions, R9). Represented values are the averages of fifty samples ± SE. Different letters indicate that the means are different by the Tukey HSD test (P ≤ 0.05).

Chlorophyll a fluorescence analysis showed that early and intermediate infested conditions were not enough to change any parameter on the three tested cultivars (data not shown). On the other hand, several parameters were affected by late infested condition. Puitá INTA-CL decreased the energy flow in photosystem II (PSII) throughout the four OJIP curve-times when comparing control and late infested plants, while both Puitá INTA-CL and BRS 7 Taim did not show any decrease in the same parameter (Fig. 2A-D). On the other hand, IRGA 423 late-infested plants increased the net rate of reaction centers closure (M_0_), reducing the energy needed to close all the reaction centers of the thylakoidal membrane (S_m_), thus showing greater efficiency in energy use, while Puitá INTA-CL and BRS 7 Taim did not present any difference compared to control plants (Fig. 2E and F). Also, Puitá INTA-CL late-infested plants reduced the fluorescence intensity at F300 (0.30 ms intensity, at the maximum point of fluorescence emission), where the plastoquinone is reduced and the reaction centers are closed, showing that this cultivar will have less energy to be used in the next phases of photosynthesis, whereas the other two cultivars present no changes in the same parameter (Fig. 2G).

**Fig. 2.**
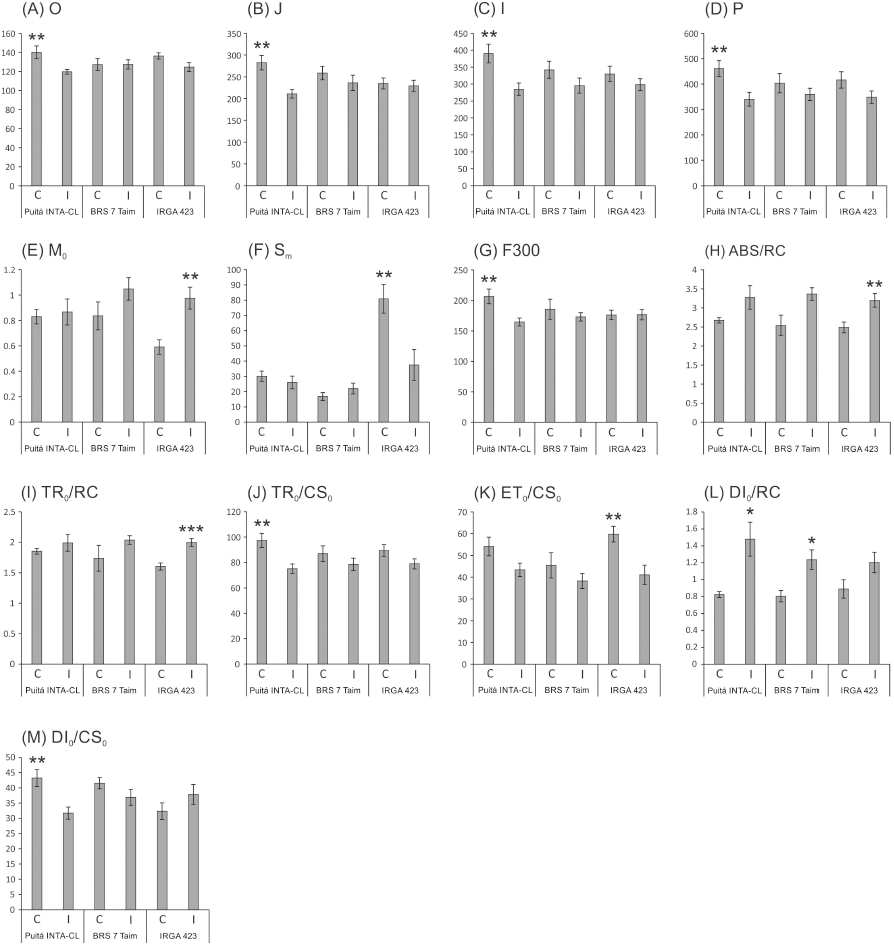
OJIP-test parameters calculated from the chlorophyll *a* fluorescence transient in control (C) and late infested (I) leaves of the three tested cultivars (Puitá INTA-CL, BRS 7 Taim, IRGA 423). (A) O; (B) J; (C) I; (D) P; (E) M_0_; (F) S_m_; (G) F300; (H) ABS/RC; (I) TR_0_ /RC; (J) TR_0_ /CS_0_; (K) ET_0_ /CS_0_; (L) DI_0_ /RC; and (M) DI_0_ /CS_0_. Represented values are the averages of ten samples ± SE. Mean values (from each cultivar) with one, two, or three asterisks are significantly different as determined by a Student’s *t* test (P ≤ 0.05, 0.01, and 0.001, respectively).

The energy absorption per reaction center (ABS/RC) is significantly increased only in infested leaves of IRGA 423 cultivar (Fig. 2H), evidencing that these plants try to obtain more energy to tolerate the herbivory stress. The light energy capture per reaction center (TR_0_/RC), which is converted in chemical energy during photosynthesis, is also increased only in infested leaves of IRGA 423 cultivar (Fig. 2I). Puitá INTA-CL was the only cultivar that decreased light energy capture per active leaf area (TR_0_/CS_0_ - Fig. 2J). The electron transport per active leaf area (ET_0_/CS_0_) decreased in infested leaves of IRGA 423 cultivar (Fig. 2K), probably decreasing ATP production as a mean of favouring defense over energy production in this cultivar. Puitá INTA-CL and BRS 7 Taim presented increased energy dissipation per reaction center (DI_0_/RC - Fig. 2L), evidencing that these two cultivars are more affected by *S. oryzae* infestation than IRGA 423, due to less efficient energy use. Puitá INTA-CL also presented a decrease in energy dissipation per active leaf area (DI_0_/CS_0_ - Fig. 2M). Based on these data, we concluded that Puitá INTA-CL and IRGA 423 show contrasting responses to *S. oryzae* infestation. Therefore, we selected these cultivars for further analysis.

The impaired chlorophyll fluorescence of Puitá INTA-CL suggests that *S. oryzae* infestation can promote an earlier senescence process on the leaves of this cultivar when compared to IRGA 423. Such hypothesis was confirmed by the higher expression of *OsSGR* gene (a senescence marker) in intermediate infested leaves of Puitá INTA-CL (Fig. 3).

**Fig. 3.**
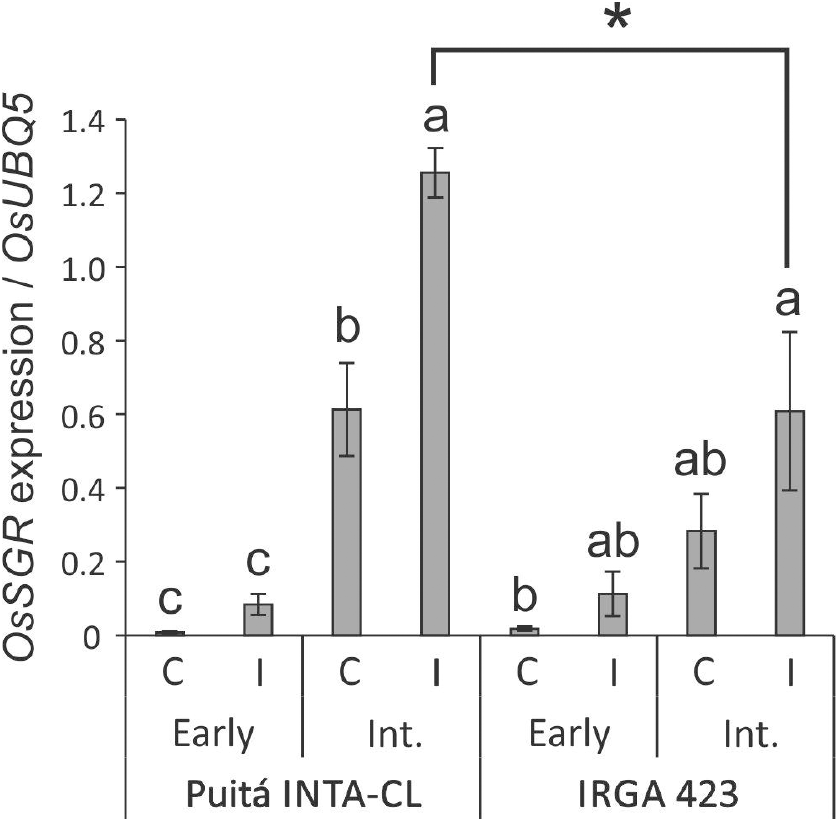
*OsSGR* expression in control and early and intermediate infested leaves of the two tested cultivars (Puitá INTA-CL and IRGA 423). Represented values are the averages of three samples ± SE. Different letters indicate that the means (from each cultivar) are different by the Tukey HSD test (P ≤ 0.05). Mean values with one asterisk are significantly different as determined by a Student’s *t* test (P ≤ 0.05).

Seeds from both cultivars were evaluated in order to verify whether *S. oryzae* infestation can decrease rice yield. As seen in Fig. 4, seeds from Puitá INTA-CL cultivar were more affected by *S. oryzae* infestation than seeds from IRGA 423, showing a decrease in the number of seeds per plant (Fig. 4A and B), percentage of full seeds (Fig. 4C), weight of 1,000 full seeds (Fig. 4D), and seed length (Fig. 4E and F), resulting in approximately 62% reduction in seed weight per plant, which is an estimative of yield lost (Fig. 5). On the other hand, seeds from IRGA 423 presented an increase in the weight of 1,000 full seeds (Fig. 4D), explained by an increase in seed length (Fig. 4F), resulting in no yield lost (Fig. 5). Based on these data, we suggest that Puitá INTA-CL is susceptible to *S. oryzae* infestation, while IRGA 423 can be considered tolerant. From now on, we will call Puitá INTA-CL and IRGA 423 as “susceptible” and “tolerant” cultivars, respectively.

**Fig. 4.**
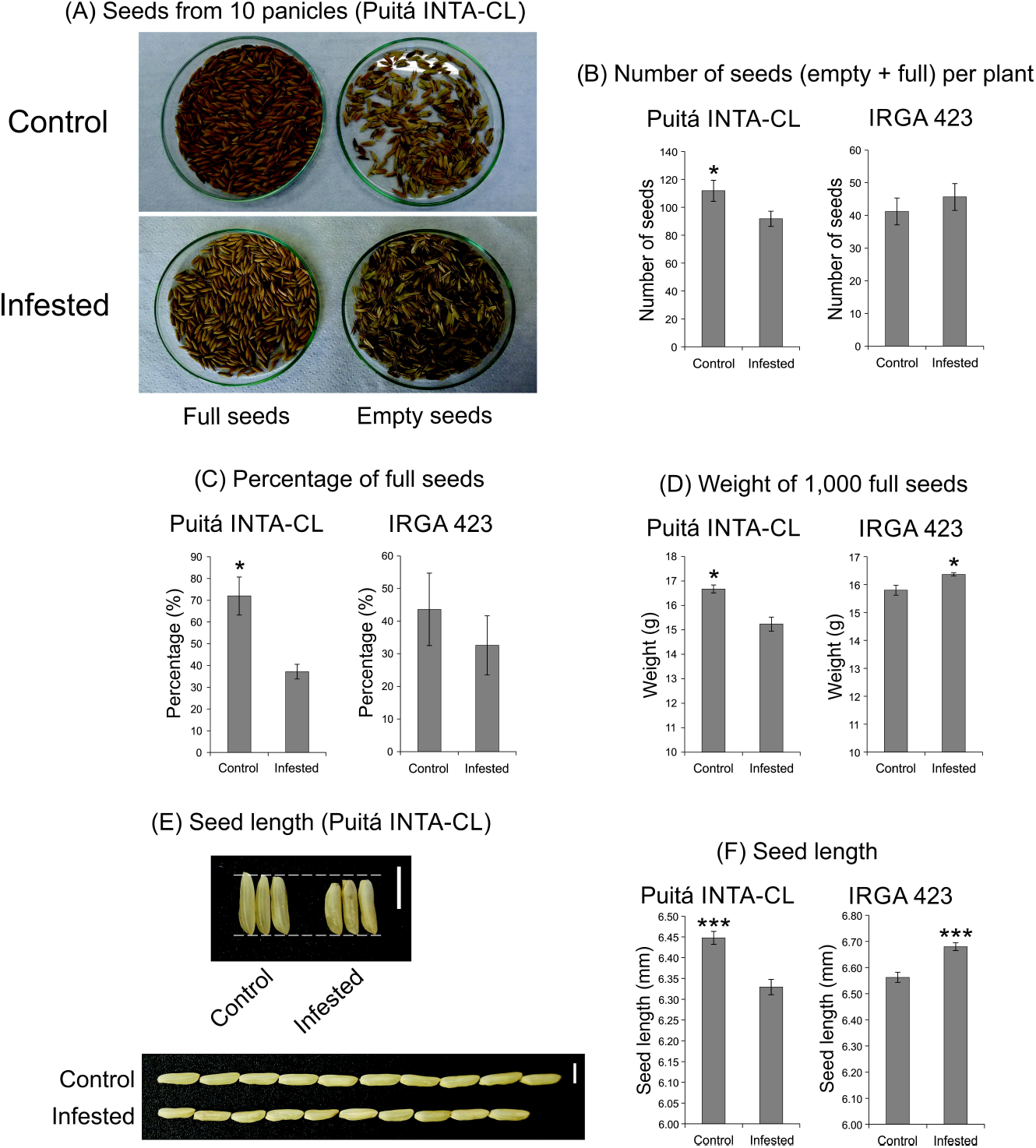
Seeds analysis of the two tested cultivars (Puitá INTA-CL and IRGA 423). (A) and (B) Number of seeds (empty + full) per plant; (C) Percentage of full seeds; (D) Weight of 1,000 full seeds; (E) and (F) Seed length (mm). Represented values are the averages of fifty samples ± SE. Mean values with one or three asterisks are significantly different as determined by a Student’s *t* test (P ≤ 0.05 and 0.001). This figure is available in colour at JXB online.

**Fig. 5.**
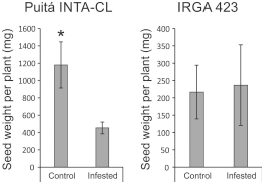
Seed weight per plant (estimative of yield) of the two tested cultivars (Puitá INTA-CL and IRGA 423). Represented values are the averages of ten samples ± SE. Mean values with one asterisk are significantly different as determined by a Student’s *t* test (P ≤ 0.05).

To verify whether *S. oryzae* infestation could differentially affect the generation of oxidative stress and the cell death level on the late infested leaves of susceptible and tolerant cultivars, we performed histochemical staining using DAB and Evans Blue. As seen in Fig. 6, leaves of the tolerant cultivar accumulates lower levels of H_2_O_2_ and less evidence of cell death (higher level of plasma membrane integrity) than the susceptible cultivar. Such low levels of oxidative stress on the leaves of tolerant cultivar could be explained, at least partially, by the higher level of phenolic compounds on the infested leaves of tolerant cultivar when compared to susceptible one (Fig. 7).

**Fig. 6.**
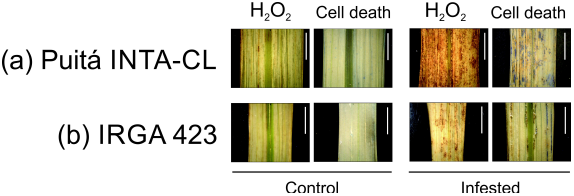
Histochemical staining assay of H_2_O_2_ and loss of plasma memebrane integrity (indicative of cell death), by diaminobenzidine (DAB) and Evans Blue, respectively, in control and late infested leaves of the two tested cultivars (Puitá INTA-CL and IRGA 423). The positive staining (detected in higher levels on infested leaves) in the photomicrographs shows as bright images (brown-color for DAB, and blue-color for Evans Blue). Bars in figures indicate 0.5 cm. This figure is available in colour at JXB online.

**Fig. 7.**
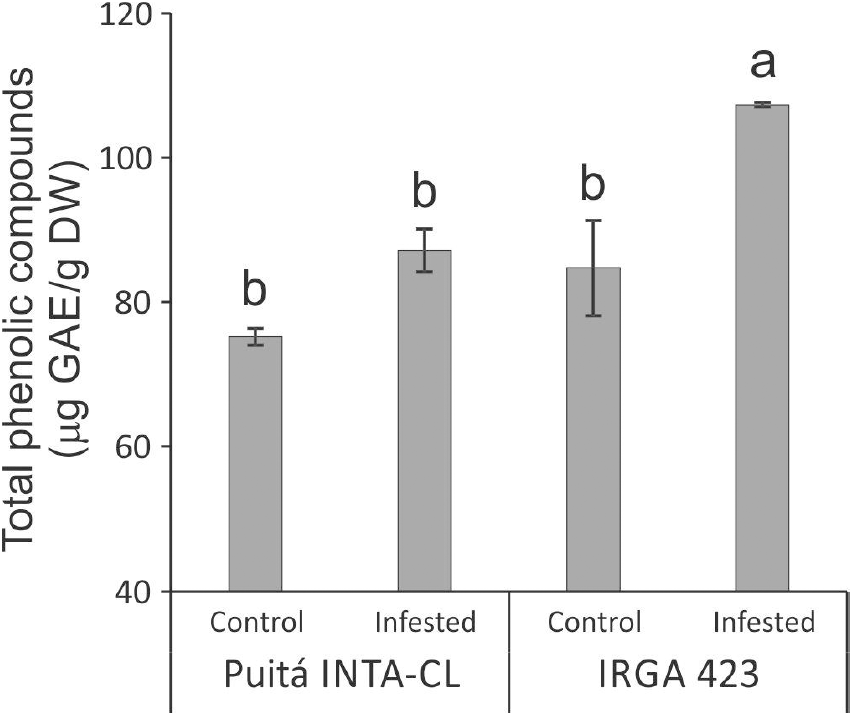
Total phenolic compounds of control and early infested leaves of the two tested cultivars (Puitá INTA-CL and IRGA 423). Represented values are the averages of three samples ± SE. Different letters indicate that the means are different by the Tukey HSD test (P ≤ 0.05). GAE: gallic acid equivalents; DW: dry weight.

We used scanning electron microscopy (SEM) to visualize the leaf surfaces of susceptible and tolerant plants during *S. oryzae* infestation. Under control condition, both cultivars presented similar levels of amorphous silica cells on the abaxial face and diminute amounts on the adaxial face (data not shown). However, under infested condition, adaxial face of tolerant IRGA 423 cultivar presents higher levels of amorphous silica cells than the susceptible Puitá INTA-CL (Fig. 8A), while similar amounts were found on the abaxial face (data not shown). Also, adaxial surface of the tolerant cultivar accumulates more SiO_2_ (major component of the amorphous silica cells) than the susceptible one (Fig. 8B).

**Fig. 8.**
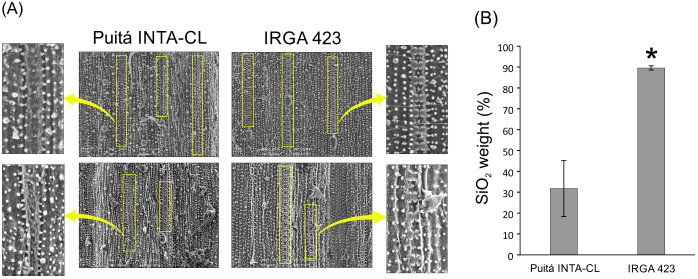
Microscope observation of amorphous silica cells and SiO_2_ quantification. (A) Scanning electron microscopy (SEM) of the infested leaf surfaces (adaxial face) from susceptible Puitá INTA-CL and tolerant IRGA 423 cultivars, highlighting the amorphous silica cells; (B) Quantification of SiO_2_ using SEM. Figures in (A) are representatives of ten analyzed leaf surfaces from each cultivar. Represented values in (B) are the averages of ten samples ± SE. Mean values with one asterisk are significantly different as determined by a Student’s *t* test (P ≤ 0.05). DW: dry weight. This figure is available in colour at JXB online.

### Overview of proteomic analysis

A crucial step in plant defense is the early perception of stress in order to respond quickly and efficiently (Rejeb *et al.,* 2014). Thus, we performed proteomic analysis of control and early infested leaves from both cultivars (susceptible and tolerant), using a Label-Free Quantitative Proteomics approach. A total of 728 proteins were identified comparing control and infested conditions in both cultivars, with 332 (45.6%) unique to or differentially abundant between cultivars. As seen in Fig. 9, comparing control and infestation leaves of susceptible cultivar, we detected 118 proteins, being 63 more abundant (and one unique) in control condition, and 54 more abundant in infested condition. We identified 217 proteins in control and infestion conditions of tolerant cultivar, being 84 more abundant (and two uniques) in control condition, and 132 more abundant (and one unique) in infested condition. When we compared both cultivars in control condition, we identified 137 proteins, with 97 more abundant (and one unique) in susceptible cultivar, and 40 more abundant in tolerant one. Comparison of both cultivars in infested condition generated 60 proteins, with 28 more abundant in susceptible cultivar, and 32 more abundant in tolerant one.

**Fig. 9.**
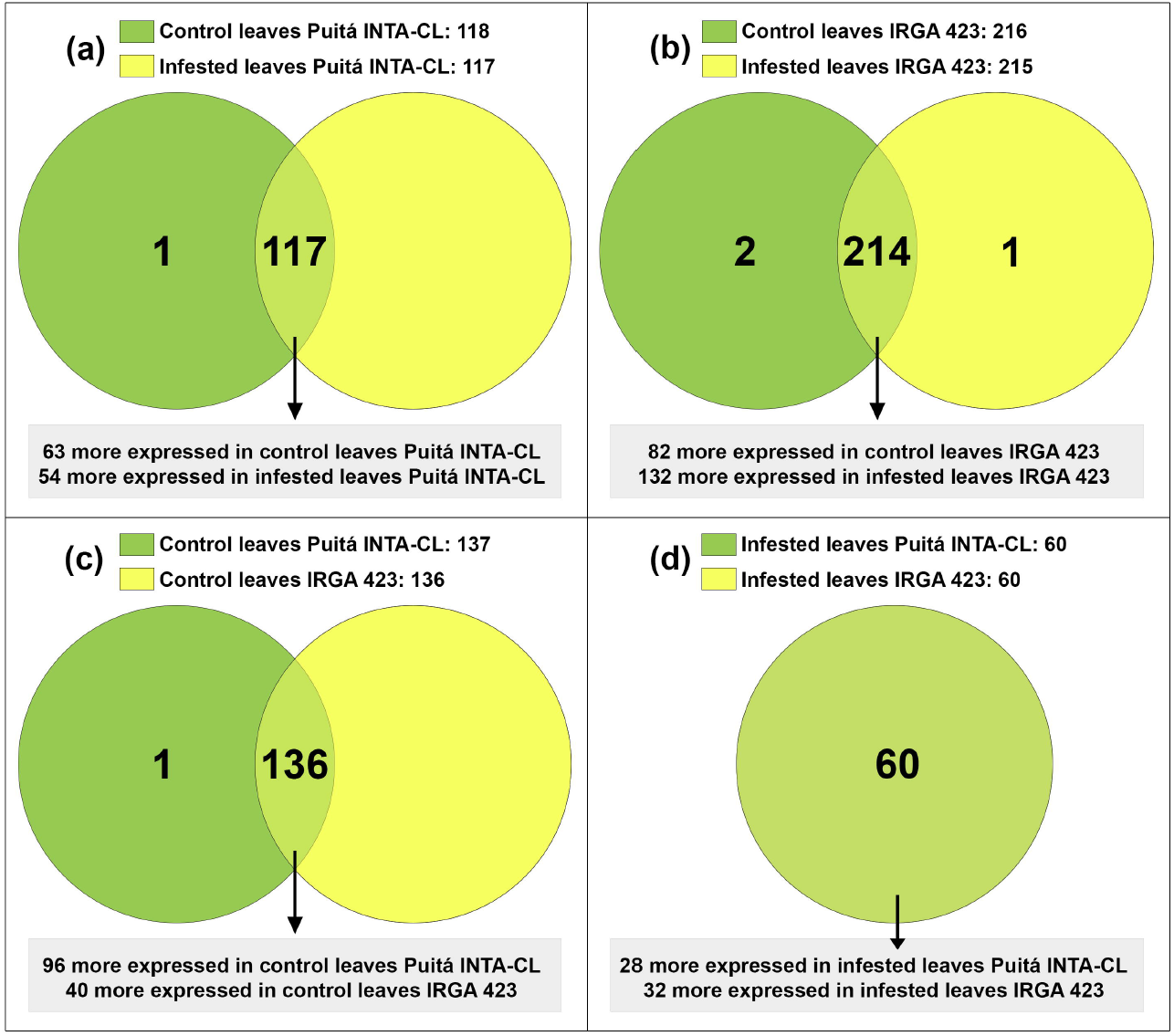
Venn diagram showing the overlap of rice proteins identified in control and early infested leaves of susceptible Puitá INTA-CL and tolerant IRGA 423 cultivars. (A) Puitá INTA-CL (control x infested); (B) IRGA 423 (control x infested); (C) control condition (Puitá INTA-CL x IRGA 423); (D) infested condition (Puitá INTA-CL x IRGA 423). In (A) and (B), dark green circles: control leaves; yellow circles: infested leaves. In (C) and (D), dark green circles: Puitá INTA-CL; yellow circles: IRGA 423. Light green means overlap in (A), (B), (C), and (D). This figure is available in colour at JXB online.

The corresponding sequence of each identified protein was submitted to NCBI BLASTp to identify specific domains, molecular functions, and protein annotations. Afterward, proteins were categorized in functional categories, according to its putative molecular function. The lists of all unique or differentially abundant proteins identified in this work are presented in Tables S2, S3, S4 and S5.

Several metabolic processes seem to be inhibited by *S. oryzae* infestation on the susceptible Puitá INTA-CL cultivar, including translation, carbohydrate metabolism and energy production (especifically glycolysis), photosynthesis and response to stress. On the other hand, the higher abundance of oxidative stress- and ATP synthesis-related proteins in infected leaves suggest an attempt to respond to *S. oryzae* infestation (Table S2). On the tolerant IRGA 423, *S. oryzae* infestation seems to be less damaging and to generate a more complex defense response. Several proteins involved with protein modification and degradation, general metabolic processes, carbohydrate metabolism and energy production (especially galactose and polysaccharide metabolism), oxidative stress, response to stress, photosynthesis, amino acid metabolism, and DNA structure maintenance were identified as more abundant in infested than in control condition. even though, some categories are still inhibited by infestation, as translation, transport, and lipid metabolism (Table S3). Surprisingly, when we compare both cultivars in control condition, the susceptible Puitá INTA-CL seems to present all the metabolic processes more active than the tolerant IRGA 423 (Table S4). However, when both cultivars are compared in infested conditions, the tolerant IRGA 423 presents higher expression of proteins related to carbohydrate metabolism/energy production and general metabolic processes than the susceptible Puitá INTA-CL, which in turn, shows increased expression of proteins related to translation and transport. Also, under infested condition, the susceptible cultivar seems to prioritize growth over defense, due to the higher expression of a gibberelin receptor, while the tolerant one seems to prioritize defense over growth, due to the higher expression of jasmonate O-methyltransferase (Table S5), a key enzyme for jasmonate-regulated plant responses.

### GO enrichment and KEGG pathways

Gene Ontology (GO) analysis provided an overview of rice molecular response to *S. oryzae* infestation in susceptible and tolerant plants. The GO annotations of all 332 differentially abundant and unique proteins identified are shown in Fig. S3 and S4. As expected, several biological processes are regulated when control and infested conditions are compared in both cultivars (control condition: Puitá INTA-CL x IRGA 423; infested condition: Puitá INTA-CL x IRGA 423), with a higher number of regulated biological processes during infestation. Two biological processes (cellular component organization and regulation of cellular processes) are more regulated on the tolerant cultivar (IRGA 423: control x infested) when compared to the susceptible one, and could be related to a more efficient plant defense (Fig. S3). The molecular function of structural constituent of ribosome is only regulated on the susceptible cultivar (Puitá INTA-CL: control x infested), and the protein binding is only regulated on the tolerant one (IRGA 423: control x infested) (Fig. S4).

To identify specific pathways affected by *S. oryzae* infestation in susceptible and tolerant rice plants, we also analyzed KEGG pathways. The following KEGG pathways (involving five or more proteins) were identified as associated with proteins differentially abundant only on the tolerant IRGA 423 cultivar (control x infested conditions): Pyruvate metabolism (11), Glyoxylate and dicarboxylate metabolism (8), Amino sugar and nucleotide sugar metabolism (8), and Methane metabolism (5). On the other hand, the following KEGG pathways (involving five or more proteins) were identified as associated with proteins differentially abundant in control condition (susceptible Puitá INTA-CL x tolerant IRGA 423): Glyoxylate and dicarboxylate metabolism (8), Citrate cycle (TCA cycle) (5), Starch and sucrose metabolism (5), Carbon fixation in photosynthetic organisms (5), Amino sugar and nucleotide sugar metabolism (5), and Frutose and mannose metabolism (5). The only pathway identified as associated with proteins differentially abundant in infested condition (susceptible Puitá INTA-CL x tolerant IRGA 423) is Gycolysis/Gluconegenesis (5), suggesting a complete different pattern of energy use employed by the cultivars in both tested conditions.

### Validation of proteomic data

The mRNA expression of three randomly selected genes (*2,3- bisphosphoglycerate-independent phosphoglycerate mutase, Hexokinase* and *Glutathione reductase*) corresponding to three proteins identified as differentially abundant during *S. oryzae* infestation was further evaluated in control and early infested leaves by RT-qPCR (Fig. 10). The proteomic profiles were confirmed for the three tested genes, even though the ratio between conditions detected at the mRNA and protein levels were different, probably due to regulation at the post-transcriptional level.

**Fig. 10.**
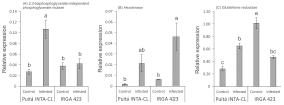
Relative expression levels (RT-qPCR, relative to *OsUBQ5* expression) of three randomly selected genes (A) *2,3-bisphosphoglycerate-independent phosphoglycerate mutase*, (B) *Hexokinase*, (C) *Glutathione reductase*, for which the encoded proteins were identified by proteomics as differentially abundant between the control and early infested leaves from Puitá INTA-CL and IRGA 423 cultivars. Represented values are the averages of three samples ± SE. Different letters indicate that the means are different by the Tukey HSD test (P ≤ 0.05).

## Discussion

In our first screening of rice responses to *S. oryzae* infestation using seven different cultivars, it was clearly shown that none of these cultivars present a classical resistance response, due to the rapid and somewhat similar infestation kinetics throughout the analyzed period (Fig. S2). Even though, physiological analysis and agronomical parameters showed that two cultivars (Puitá INTA-CL and IRGA 423) present different responses to *S. oryzae* infestation (Fig. 1-8), being considered susceptible and tolerant, respectively. In fact, there is not a consensus about the requirement for a trait be considered as a plant defense mechanism (Karban, 2011; Poelman, 2015), and most plant defenses are still characterized by proximate variables such as herbivore performance or plant damage (Wetzel *et al.,* 2016; Erb, 2018). However, plant defenses can be surely defined as traits that reduce the negative impact of herbivores on plant reproductive success or that increase plant fitness (Erb, 2018). According to Peterson *et al.* (2017), tolerance mechanisms allow the plants to withstand pest injury and produce acceptable yields, maintaining the fitness under stressful conditions. For this reason, the fact that IRGA 423 did not decrease the yield under infested condition (compared to 62% in Puitá INTA-CL) was the main characteristic that encouraged us to define IRGA 423 as tolerant to *S. oryzae* infestation.

Chlorophyll fluorescence is a non-invasive tool commonly used for determining the behavior of the photosynthetic apparatus of control and abiotic stressed plants (Gururani *et al.,* 2015; Rapacz *et al.,* 2015). Our group previously detected a reduction in Pi_ABS_, S_m_, and N parameters (related to the donor and acceptor sides of PSII) in rice leaves early infested by *S. oryzae* in IRGA 424 cultivar (Buffon *et al.,* 2016). However, to the best of our knowledge, this is the first work that uses this type of photosynthetic analysis to differentiate plant susceptibility and tolerance to a biotic stress. Several parameters related to chlorophyll *a* fluorescence were affected in the susceptible Puitá INTA-CL cultivar during infestation, showing a worse photosynthetic performance than IRGA 423 (Fig. 2). Puitá INTA-CL increased its energy dissipation per reaction center (DI_0_/RC - Fig. 2L) and decreased energy flow in PSII at the four OJIP-curve/times (Fig. 2A-D), fluorescence intensity at F300 (Fig. 2G), energy dissipation per active leaf area (DI_0_/CS_0_ - Fig. 2M), and light energy capture per active leaf area (TR_0_/CS_0_ - Fig. 2J), the last one probably linked to enhanced cell death in their leaves (Fig. 6). Zhang *et al.* (2013) demonstrated that excess Ca^2+^ increased the toxicity of Hg^2+^ to PSII of cyanobacterium *Synechocystis* sp. through the increased of energy flux dissipation per reaction center (DI_0_/RC), leading to dysfunction of PSII. Rapacz *et al.* (2015) showed that DI_0_/RC parameter increase with increasing levels of PSII damage in wheat under low temperature. The I and P fluorescence intensities on the OJIP induction curve and the F300 parameter also decrease in wheat plants exposed to Pb stress (Kalaji and Loboda, 2007). Interestingly, tall fescue *Festuca arundinacea* leaves show a great decrease at all steps of OJIP and F300 parameters in response to high-temperature stress, but pre-acclimation treatment inhibit such declines (Hu *et al.,* 2015). Intriguingly, Rapacz *et al.* (2015) suggest that DI_0_/CS_0_ values increase with increasing levels of PSII damage, which is the opposite to what we found (Fig. 2M). More studies are needed to clarify the impact of DI_0_/CS_0_ in rice photosynthetic performance. In common wheat, TR_0_/CS_0_ values correlate well with plant survival after freezing, being an excellent indicator for prediction of winter field survival or estimation of freezing tolerance (Rapacz *et al.,* 2015). According to Gururani *et al.* (2015) and Rapacz *et al.* (2015), the OJIP test is a reliable indicator of cold tolerance in the turfgrass *Zoysia japonica* and freezing tolerance in wheat, respectively. Therefore, we suggest for the first time that rice tolerance to *S. oryzae* (and probably to other herbivores) can also be estimated by OJIP test. According to Peterson *et al.* (2017), increased net photosynthetic rate after herbivory is one of the general physiological mechanisms involved in plant tolerance. The differences in photosynthetic performance presented by the susceptible Puitá INTA-CL and tolerant IRGA 423 cultivars are supported by decreased and increased numbers of photosynthesis-related proteins detected in response to *S. oryzae* infestation, respectively (Tables S2 and S3).

As a result of impaired chlorophyll *a* fluorescence in infested leaves of Puitá INTA-CL, an earlier senescence process was established in their leaves upon *S. oryzae* infestation (Fig. 3). Leaf senescence is a natural and important developmental process, responsible for great part of the nitrogen mobilized to the seeds. Late senescence, which means a prolonged and maximum period of photosynthetic activity, should lead to higher yields (Jagadish *et al.,* 2015; Diaz-Mendoza *et al.,* 2016). However, senescence processes are also closely linked to stress conditions, which commonly anticipate this process (Wojciechowska *et al.,* 2017). Therefore, manipulation of senescence events can be a rationale way to obtain higher yield and quality of grains (Egli, 2011; Jagadish *et al.,* 2015). We believe that the late senescence process detected on the leaves of tolerant IRGA 423 cultivar is at least partially responsible for the better seed characteristics presented by this cultivar under *S. oryzae* infestation (Fig. 4), including the absence of yield loss. Yet, infested leaves of susceptible Puitá INTA-CL cultivar express atATG18b protein (Table S5), which is required for the formation of autophagosomes during nutrient stress and senescence in Arabidopsis (Xiong *et al.,* 2005).

Even though we detected a lower level of H_2_O_2_ accumulation in infested leaves of the tolerant IRGA 423 cultivar (Fig. 6), we were not able to find a clear difference in oxidative stress-related proteins identified on the both cultivars under infested condition (Tables S2, S3, and S5), except a peroxidase protein 2.6-fold more abundant in tolerant IRGA 423 infested leaves (Supplmentary Table 5). However, infested leaves of the tolerant cultivar accumulate higher levels of phenolic compounds than the susceptible one (Fig. 7). In plants, it is well established that phenolics can act as antioxidants by donating electrons to guaiacol-type peroxidases for the detoxification of H_2_O_2_ produced under different stress conditions, including biotic ones (Sakihama *et al.,* 2002; Shalaby and Horwitz, 2015; Hung, 2016). Also, many structurally different phenolics rapidly accumulate to higher levels as components of an induced defense arsenal against herbivore attack (Gaquerel *et al.,* 2014; Karabourniotis *et al.,* 2014). For example, the larval development of the pea aphid is longer, the reproduction period is shorter, the fecundity is decreased, and the aphid population is reduced on alfalfa lines containing high levels of phenolics (GoŁawska and Łukasik, 2009). Therefore, we believe that part of the tolerance mechanism to *S. oryzae* infestation by IRGA 423 cultivar is due to phenolics accumulation in their leaves. Interestingly, the accumulation of phenolic compounds along with enhancement of phenylpropanoid metabolism has been observed under different environmental stress conditions (Michalak, 2006). Phenylpropanoid metabolic pathway synthesize flavonoids, which have many diverse functions, including plant responses to stress conditions (as cold and drought - Schulz *et al.,* 2016; Shojaie *et al.,* 2016) and defense (Buer *et al.,* 2010), functioning as powerful antioxidants. Under infested condition, we detected flavonol-3-O-glycoside-7-O-glucosyltransferase 1 protein, involved with flavonol biosynthesis (Sun *et al.,* 2016), 3.6-fold more abundant on the tolerant IRGA 423 cultivar than the susceptible one (Table S5), suggesting that lavonoids can also contribute to tolerate *S. oryzae* infestation.

The beneficial effects of silicon (Si) on plant resistance against biotic stresses, including insect herbivory, have been well documented in rice plants, showing positive correlations between increased Si content and enhanced insect resistance (Sidhu *et al.,* 2013; Ye *et al.,* 2013; Han *et al.,* 2016). We detected higher number of amorphous silica cells and higher accumulation of SiO_2_ in infested leaves of the tolerant IRGA 423 cultivar when compared to the susceptible one (Fig. 8). Based on these data, we strongly suggest that enhanced Si levels can also contribute to the more effective defense of IRGA 423 cultivar against *S. oryzae* mite infestation. According to Ye *et al.* (2013), there is a strong interaction between Si and JA in rice defense against insect herbivores involving priming of JA-mediated defense responses by Si and the promotion of Si accumulation by JA. This is reinforced by the higher expression of jasmonate O-methyltransferase protein on the infested leaves of tolerant IRGA 423 cultivar (Table S5). Such enzyme catalyzes the methylation of JA into Me-JA, which controls plant defenses against herbivore attack (Qi *et al.,* 2016; Huang *et al.,* 2017). On the other hand, we detected higher expression of a gibberelin (GA) receptor protein on the infested leaves of susceptible Puitá INTA-CL cultivar (Table S5). GA regulates many essential plant developmental processes, including growth (Hou *et al.,* 2013). As GA and JA antagonize each other in regulating plant growth and defense (Chaiwanon *et al.,* 2016;Ning *et al.,* 2017), we suggest that under infested condition, the susceptible Puitá INTA-CL cultivar prioritize growth over defense, while the tolerant IRGA 423 prioritize defense over growth, and this difference might contribute to the *S. oryzae* susceptiblity or tolerance.

## Conclusion

This is the first report evaluating the defense responses of two contrasting rice cultivars to *Schizotetranychus oryzae* infestation. Physiological analyses showed that the tolerant IRGA 423 cultivar under infested condition presents better photosynthetic performance, later leaf senescence period, less affected seeds and lower yield loss, lower levels of H_2_O_2_ accumulation and cell death, higher levels of phenolic compounds (and probably flavonoids), higher level of SiO_2_ concentration in leaves, and higher number of amorphous silica cells on the leaf surface than the susceptible Puitá INTA-CL cultivar. Proteomic analysis showed that the tolerant IRGA 423 cultivar under infested condition presents a more complex and efficient response to *S. oryzae* infestation, with carbohydrate metabolism/energy production, general metabolic processes, and JA biosynthesis more active than the susceptible Puitá INTA-CL cultivar. The model in Fig. 11 summarize the rice tolerance mechanisms employed by IRGA 423 cultivar. Altogether, these findings are helpful to reveal the different molecular mechanisms involved in the rice response to *S. oryzae* infestation, and could be used in future breeding programs or genetic engeneering attempts aiming to increase mite tolerance in rice plants.

**Fig. 11.**
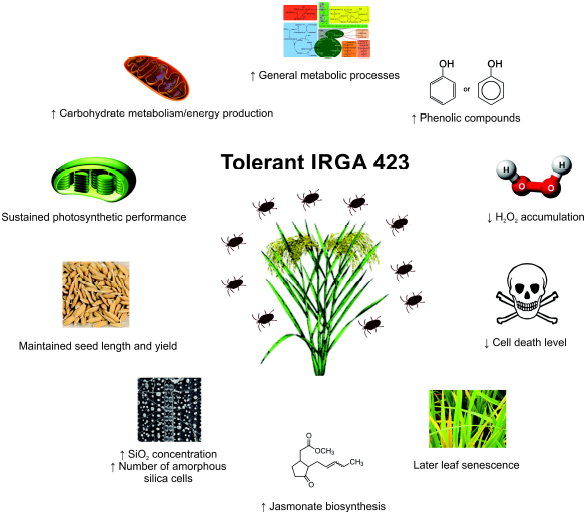
Rice mechanisms employed by IRGA 423 cultivar to tolerate *Schizotetranychus oryzae* infestation. This figure is available in colour at JXB online.

## Supplementary Data

**Table S1.** Gene-specific PCR primers used for RT-qPCR analyses.

**Table S2.** Differentially abundant proteins in susceptible Puitá INTA-CL cultivar (control x infested condition).

**Table S3.** Differentially abundant proteins in tolerant IRGA 423 cultivar (control x infested condition).

**Table S4.** Differentially abundant proteins in control condition (susceptible Puitá INTA-CL x tolerant IRGA 423).

**Table S5.** Differentially abundant proteins in infested condition (susceptible Puitá INTA-CL x tolerant IRGA 423).

**Figure S1.** Classification of infestation levels according to visual characteristics of leaves.

**Figure S2.** Pattern of infestation kinetics after five weeks in the seven tested cultivars.

**Figure S3.** Gene ontology annotation. Biological processes of differentially abundant and unique proteins obtained from control and early infested leaves from susceptible Puitá INTA-CL and tolerant IRGA 423 cultivars.

**Figure S4.** Gene ontology annotation. Molecular functions of differentially abundant and unique proteins obtained from control and early infested leaves from susceptible Puitá INTA-CL and tolerant IRGA 423 cultivars.

## Acknowledgements

This research was supported by University of Taquari Valley - UNIVATES. The authors thank Instituto Rio-Grandense do Arroz (IRGA) for technical support, and JosÉ Rafael Wanderley Benicio and Rafael Spiekermann for the use of stereomicroscope.

## Abbreviations

EI: early infestation
II: intermediate infestation
JA: jasmonate
LI: late infestation
PSII: photosystem II
SEM: scanning electron microscopy

